# Innate IFNγ is essential for systemic *Chlamydia* control while CD4 T cell-dependent IFNγ production is highly redundant in the female reproductive tract

**DOI:** 10.1101/2020.08.26.269480

**Authors:** Miguel A.B. Mercado, Wuying Du, Priyangi A. Malaviarachchi, Jessica I. Gann, Lin-Xi Li

## Abstract

Protective immunity to the obligate intracellular bacterium *Chlamydia* is thought to rely on CD4 T cell-dependent IFNγ production. Nevertheless, whether IFNγ is produced by other cellular source during *Chlamydia* infection and how CD4 T cell-dependent and -independent IFNγ contribute differently to host resistance has not been carefully evaluated. In this study, we dissect the requirements of IFNγ produced by innate immune cells and CD4 T cells for resolution of *Chlamydia muridarum* female reproductive tract (FRT) infection. After *C. muridarum* intravaginal inoculation, IFNγ-deficient and T cell-deficient mice exhibited opposite phenotypes for survival and bacterial shedding at the FRT mucosa, demonstrating the distinct requirements for IFNγ and CD4 T cells in host defense against *Chlamydia*. In *Rag*-deficient mice, IFNγ produced by innate lymphocytes (ILCs) accounted for early bacterial containment and prolonged survival in the absence of adaptive immunity. Although group I ILCs are potent IFNγ producers, we found that mature NK cells and ILC1 were not the sole source for innate IFNγ in response to *Chlamydia*. T cell adoptive transfer experiments revealed that WT and IFNγ-deficient CD4 T cells were equally capable of mediating effective bacterial killing in the FRT during the early stage of *Chlamydia* infection. Together, our results revealed that innate IFNγ is essential for preventing systemic *Chlamydia* dissemination, whereas IFNγ produced by CD4 T cells is largely dispensable at the FRT mucosa.

## Introduction

*Chlamydia trachomatis* is the obligate intracellular bacterium that causes the most prevalent sexually transmitted infection worldwide. The prevalence of Chlamydia is partially attributed to its nature of being a “silent infection”, as most women infected with *C. trachomatis* are asymptomatic and can resolve the infection spotanously (1). Unfortunately, undiagnosed infections not only facilitate silent disease transmissions, but can also lead to severe adverse effects such as pelvic inflammatory disease, ectopic pregnancy and infertility (2, 3). There is no licensed human Chlamydia vaccine available at present, partially owing to the lack of complete understanding of protective immune mechanisms (4–6).

IFNγ produced by CD4 Th1 cells promotes macrophage activation to eliminate intracellular pathogens within these professional phagocytes. This defense mechanism operates efficiently against bacterial pathogens that exhibit marked infection tropism for macrophages (7, 8). For *Chlamydia*, infection is largely restricted to epithelial cells at barrier tissues such as the eyes, lungs and the reproductive tract. As such, it is reasonable to speculate that protective immunity to *Chlamydia* would be more complex than Th1-dependent IFNγ production (9).

The mouse models of *Chlamydia* female reproductive tract infection provide an invaluable tool for studying host immunity to *Chlamydia* infection. Research in the field has established a predominant role for CD4 T cells and antibody in host resistance and vaccine afforded protection against *Chlamydia* (4, 10–12). After *Chlamydia muridarum* intravaginal infection, WT B6 mice exhibit a self-limiting infection that resolves within 4-5 weeks. Similar courses of infection were also reported in mice lacking B cells, CD8 T cells and γδT cells, demonstrating the redundant roles of these cells in *C. muridarum* clearance from the female reproductive tract (FRT) (13–15).

In contrast, nude mice and mice lacking MHC class II-restricted CD4 T cells shed persistent high levels of bacteria from the FRT without showing obvious signs of wasting (15–17). While a large body of evidence points to a role for Th1, but not other Th lineage cells in protective immunity against *Chlamydia* (14, 18–23). it remains perplexing that IFNγ-deficient mice are capable of eliminating >99% of *C. muridarum* from the FRT, albeit suffering from multiorgan disseminated infections (14, 24).

The distinct phenotypes of CD4 T cell-deficient and IFNγ-deficient mice after *C. muridarum* infection prompted us to investigate the definitive contributions of CD4 T cells-dependent and - independent IFNγ production in host defense against *Chlamydia*. We hypothesize that IFNγ produced during innate immune response prevents lethal disseminated infection, while CD4-dependent IFNγ production is largely dispensable at the FRT mucosa. We tested these hypotheses using loss- and gain-of-function approaches in gene-deficient mouse models in which contributions of innate and adaptive IFNγ production can be dissected separately.

## Materials and Methods

### Mice

C57BL/6 (B6), IFNγ^−/−^ (B6.129S7-*Ifng*_*tm1Ts*_/J), *Rag1*^−/−^ (B6.129S7-*Rag1*_*tm1Mom*_/J), TCRβ^−/−^ (B6. 129P2-*Tcrb*_*tm1Mom*_/J) mice were purchased from The Jackson Laboratory. *Rag2*^−/−^*γc*^−/−^ mice were purchase from Taconic Biosciences. All mice used for experiments were 6-16 weeks old, unless otherwise noted. Mice were maintained under SPF conditions and all mouse experiments were approved by University of Arkansas for Medical Sciences (UAMS) Institutional Animal Care and Use Committee (IACUC).

### Bacteria

*Chlamydia muridarum* strain Nigg II was originally purchased from ATCC (VR-123; Manassas, VA). The organism was propagated in HeLa 229 cells. Elementary bodies (EBs) were purified by renografin discontinuous density gradient centrifugation, aliquoted and stored at −80°C until use (25). EBs were titrated on HeLa 229 cells as described below.

### Infection and bacteria enumeration

Mice were synchronized for estrous by subcutaneous injection of 2.5 mg Depo-provera (Greenstone, NJ), 5-7 days prior to intravaginal infection. For intravaginal infection, 1×10^5^ *C. muridarum* in SPG buffer were deposited directly into the vaginal vault using a pipet tip. To enumerate bacterial shedding from the FRT, vaginal swabs were collected, suspended in SPG buffer and disrupted with glass beads. Inclusion forming units (IFUs) were determined by plating serial dilutions of swab samples on HeLa 229 cells, staining with anti-MOMP mAb and counting under microscope. To enumerate bacteria burden within tissues, intraperitoneal (IP) wash was collected in SPG buffer, spleen, kidney, lung and gall bladder were homogenized in SPG buffer.

Tissue homogenates were disrupted with glass beads, centrifuged at 500g for 10 minutes, and supernatants collected and serial dilutions plated on HeLa 229 cells for IFU counts.

### Ab-mediated depletion

IFNγ in vivo depletion was performed by IP injection of 0.25 mg anti-IFNγ (XMG1.2, BioXcell) on days −1, 1, and every 3 days thereafter post infection. NK cells were depleted by IP injection of 0.3 mg anti-NK1.1 (purified mAb from PK136 hybridoma, gift from Richard Morrison, UAMS) on days −3, −1, 1, and every 3 days thereafter post infection.

### CD4 T cell adoptive transfer

Total CD4 T cells from donor mice were purified from spleens using STEMCELL EasySep CD4 T Cell Isolation Kit according to manufacturer’s instruction (STEMCELL Technologies). Depending on individual experiment, five to twenty million purified CD4 T cells were transferred intravenously into recipient mice via the tail vein.

### IFNγ secretion assay and flow cytometry

IFNγ secretion assay was conducted according to manufacturer’s instruction (Miltenyi, Mouse IFNγ Secretion Assay Detection Kit). Briefly, spleens were harvested and single cell suspensions prepared in RPMI with 5% FCS. Cells were labeled with IFNγ Catch Reagents for 5 min on ice and incubation for 45 min at 37°C. IFNγ-producing cells were stained with IFNγ Detection Antibody in conjunction with cell surface markers (listed below) and analyzed on an LSRFortessa flow cytometer (BD Biosciences). Antibodies used included CD3e (145-2C11), CD11b (M1/70) and NK1.1 (PK136) (BioLegend). Data were analyzed using FlowJo software (Tree Star).

### Statistical analysis

Statistical analysis was performed with GraphPad Prism 8. Unpaired *t* test was used for normally distributed continuous-variable comparisons; Mann-Whitney U test was used for nonparametric comparisons. The log-rank Mantel-Cox test was used for survival curves.

## Results

### IFNγ-deficient and T cell-deficient mice exhibit opposite phenotypes after Chlamydia muridarum intravaginal infection

In order to directly compare the phenotypes of IFNγ-deficient and T cell-deficient mice, we infected IFNγ^−/−^ and TCRβ^−/−^ mice intravaginally with *C. muridarum* strain Nigg II and compared survival and bacterial shedding during primary infections. Consistent with previous findings (14), one hundred percent of TCRβ^−/−^ mice survived the infection with no obvious sign of wasting disease (Fig. 1A). Meanwhile, these mice manifested high-grade, persistent bacterial shedding with an average of >10^5^ bacteria recovered from the FRT for at least 60 days (Fig.1A and 1B). In contrast, over 90% of IFNγ^−/−^ mice succumbed to infection between days 15 and 47 after infection (Fig. 1A). Notably, IFNγ^−/−^ mice that survived by day 35 post infection exhibited ~10,000-fold reduction in FRT bacterial burden compared to day 7 (Fig. 1B). These opposite phenotypes of T cell-deficient and IFNγ-deficient mice suggest that distinct host defense mechanisms are involved in IFNγ− and T cell-dependent *Chlamydia* containment in systemic versus mucosa tissues. Importantly, a non-αβT cell source of IFNγ in TCRβ^−/−^ mice must have contributed to systemic *Chlamydia* control for their long-term survival.

**Fig. 1.**
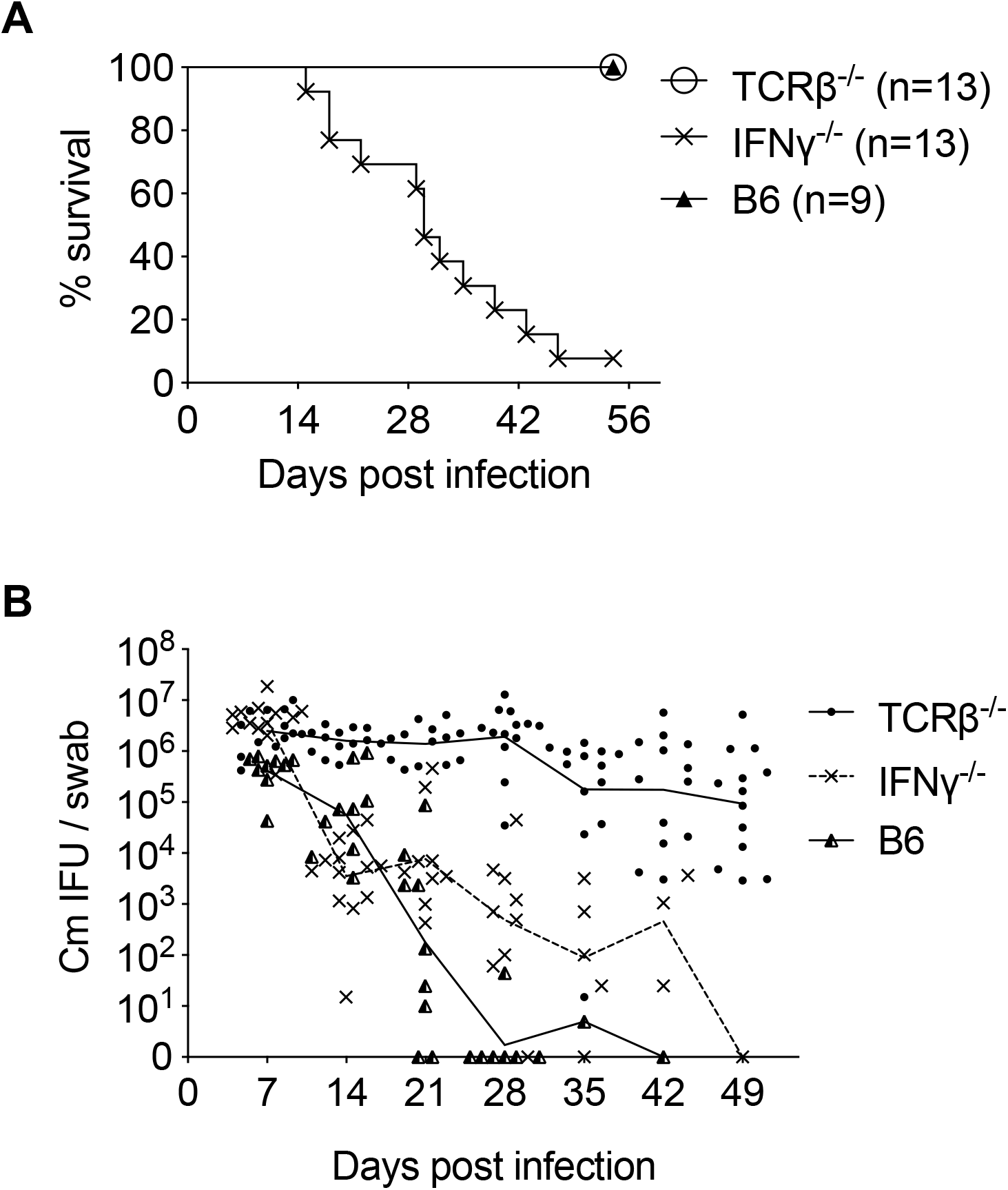
IFNγ-deficient mice and αβT cell-deficient mice exhibit opposite phenotypes after *Chlamydia muridarum* intravaginal infection. B6, IFNγ^−/−^ and TCRβ^−/−^ mice were infected intravaginally with 1×10^5^ *C. muridarum*. (A) Survival and (B) bacterial shedding from the lower female reproductive tract (FRT) were monitored by vaginal swabs. Data shown are combined results of three independent experiments of 9-13 mice per group. Each data point represents an individual mouse. Lines represent mean log_10_-transformed values.

### ILCs are essential for systemic bacterial control and prolonged survival of Rag1^−/−^ mice

The non-lethal phenotype of TCRβ^−/−^ mice led us to hypothesize that innate lymphocytes (ILCs) are essential for preventing lethal *Chlamydia* dissemination. To test this hypothesis, we infected *Rag1*^−/−^ and *Rag2*^−/−^*γc*^−/−^ mice intravaginally with *C. muridarum* and monitored survival. *Rag2*^−/−^*γc*^−/−^ mice quickly succumbed to infection with a median survival time of 12 days. This is significantly shorter compared to the average 30 days of survival in *Rag1*^−/−^ mice (Fig. 2A). Fast weight loss was observed in *Rag2*^−/−^*γc*^−/−^ mice starting from day 5 post infection, but not in *Rag1*^−/−^ mice (Fig. 2B). By the time *Rag2*^−/−^*γc*^−/−^ mice were moribund (10-13 dpi), we detected widespread bacteria dissemination in these mice with high bacterial burdens in systemic tissues including spleen, kidney, lung, ball bladder, peritoneal cavity, as well as vaginal and rectal mucosa (Fig. 1C). Notably, while all systemic tissues of *Rag2*^−/−^*γc*^−/−^ mice exhibit 1-4 logs higher bacteremia than *Rag1*^−/−^ mice, *Chlamydia* burdens at the FRT mucosa were not significantly different between the two strains (Fig. 1C). These results demonstrate that ILCs are essential for systemic *Chlamydia* control, but are redundant for host resistance to *Chlamydia* at the FRT mucosa in the absence of adaptive immunity.

**Fig. 2.**
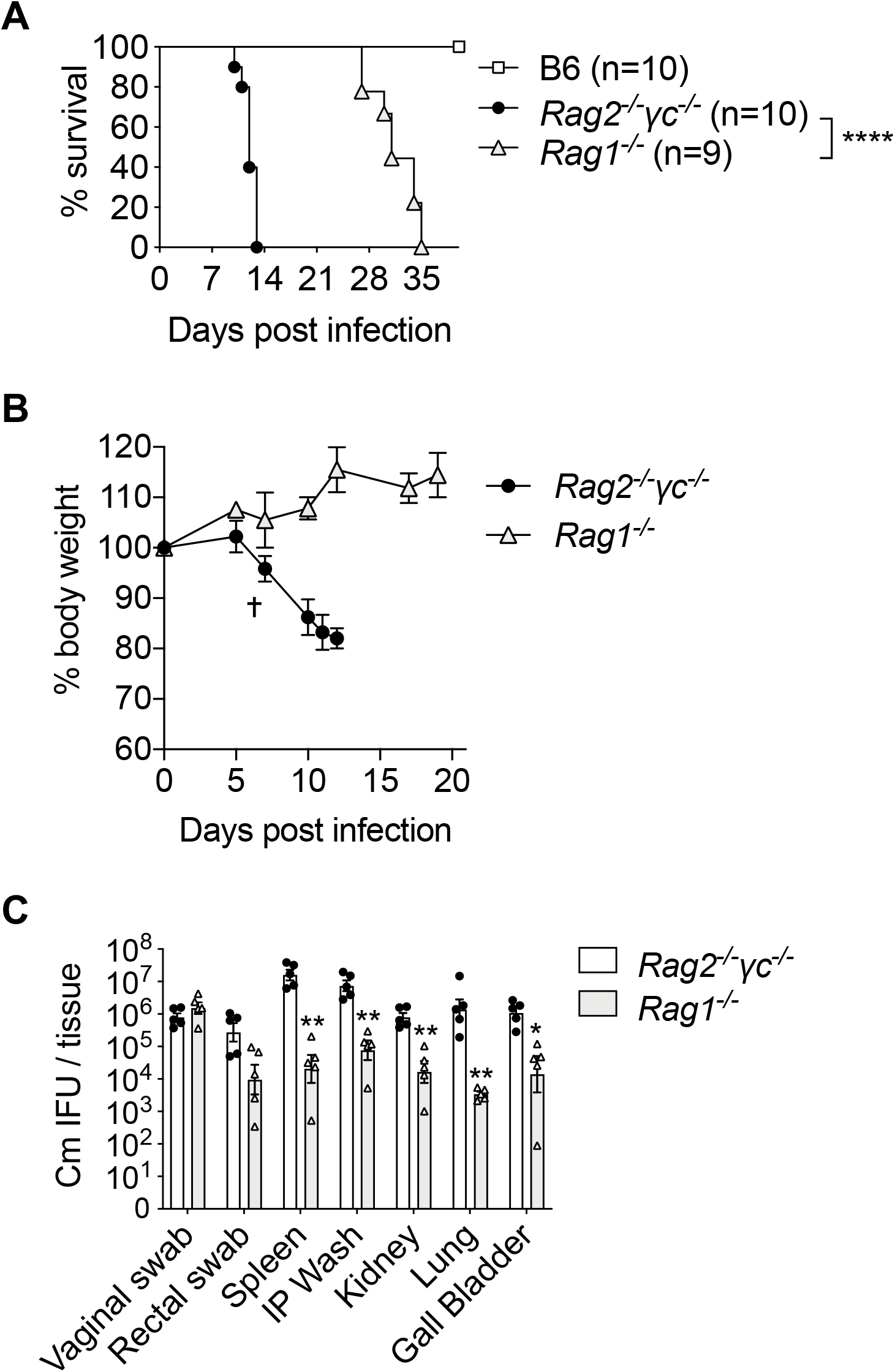
Innate lymphocytes (ILCs) are essential for systemic bacterial control and prolonged survival of *Rag1*-deficient mice. B6, *Rag1*^−/−^ and *Rag2*^−/−^*γc*^−/−^ mice were infected intravaginally with 1×10^5^ *C. muridarum*. (A) Survival and (B) body weight were monitored. (C) Bacterial burdens in systemic organs were determined at 12 days post infection. Data shown are (A) combined results of two independent experiments of 9-10 mice per group or (B-C) representative results of two independent experiments with 3-5 mice per group in each experiment. Each data point represents an individual mouse. Error bars represent the mean ± SEM; *p < 0.05; **p < 0.01; ****p < 0.0001.

### Innate IFNγ production, partially by NK cells and ILC1s, is essential for early control of Chlamydia dissemination

We next investigate whether IFNγ production by ILCs, in particular the group 1 ILCs including NK cells and ILC1s, accounts for the key innate effector mechanism for early control of *C. muridarum* systemic dissemination. To do this, we depleted either IFNγ or NK1.1^+^ cells from *Rag1*^−/−^ mice and monitored survival. As shown in Fig. 3A, both IFNγ− and NK1.1-depleted groups displayed accelerated death compared to PBS treated group, with average survival time of 16.5 and 21 days, respectively. IFNγ secretion by mature group 1 ILCs (CD11b^+^NK1.1^+^) were readily detectable in the spleens of *Rag1*^−/−^ mice throughout the course of infection, but not evident in WT B6 controls (Fig. 3B and 3C). Unexpectedly, circulating IFNγ levels in *Rag1*^−/−^ mice were not significantly affected by NK1.1 depletion at both days 7 and 14 post infection (Fig. 3D). Together, these findings led us to conclude that both innate IFNγ and group 1 ILCs are essential for early containment of *Chlamydia* dissemination. While NK1.1^+^ group 1 ILCs are potent IFNγ producers, they do not seem to be the only source for IFNγ derived from innate immune responses following *C. muridarum* intravaginal infection.

**Fig. 3.**
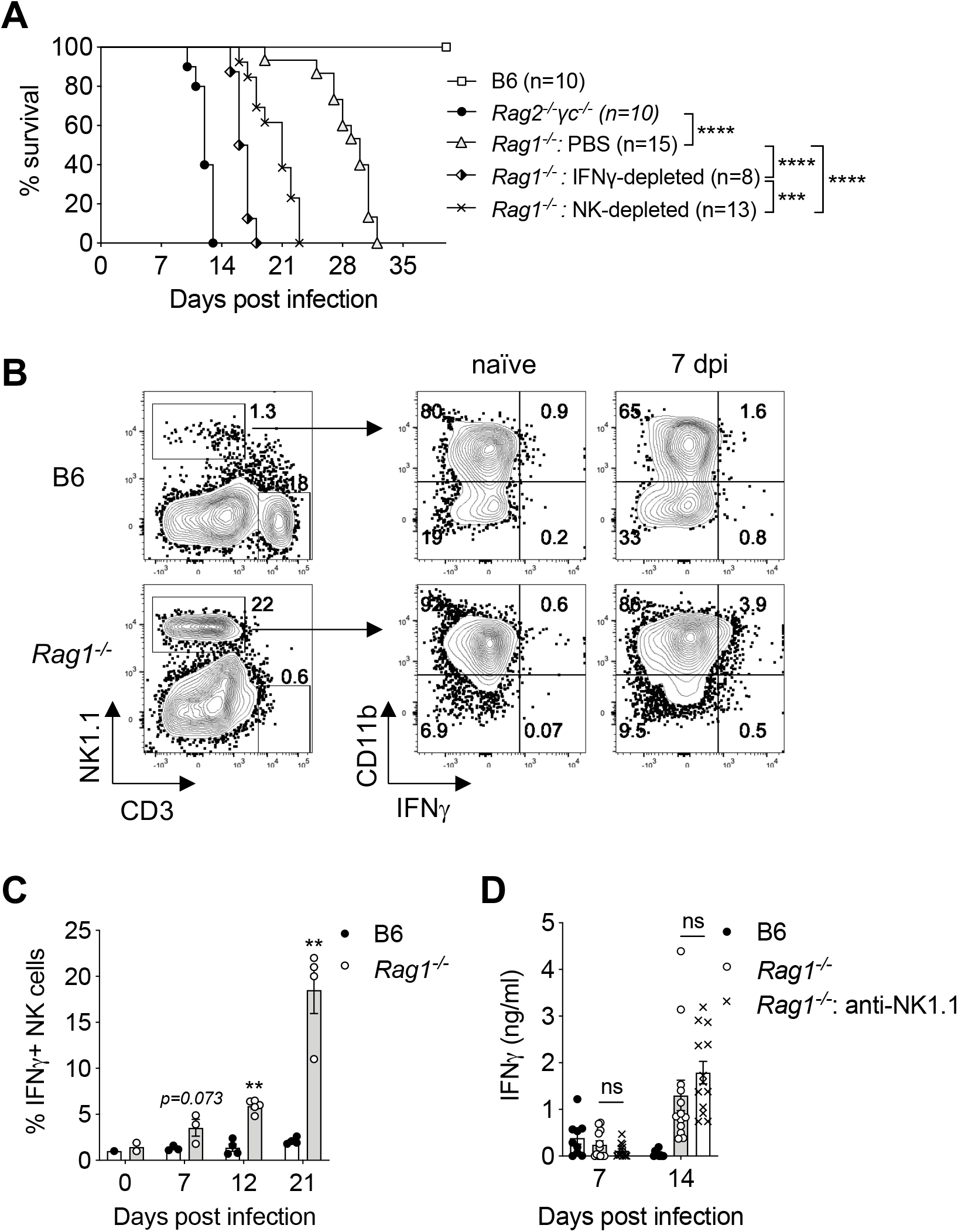
Innate IFNγ and group 1 ILCs contribute to host resistance to lethal *Chlamydia* dissemination in *Rag1*^−/−^ mice. *Rag1*^−/−^ mice were infected intravaginally with 1×10^5^ *C. muridarum*. Groups of mice were treated with either anti-IFNγ or anti-NK1.1 depleting Abs throughout the course of infection. (A) Survival was monitored. (B) Representative flow cytometry plots showing IFNγ secretion by CD11b^+^NK1.1^+^ cells detected by IFNγ secretion assay. (C) Percentages of IFNγ-producing CD11b^+^NK1.1^+^ cells were quantified based on flow cytometry analysis in (B). (D) Serum IFNγ level on days 7 and 14 after infection as measured by IFNγ cytokine ELISA. Data shown in (A), (C), and (D) are combined results of at least two independent experiments with 3-5 mice per group in each experiment. Each data point in (C) and (D) represents an individual mouse. Error bars represent the mean ± SEM; *p < 0.05; **p < 0.01; ***p < 0.001; ****p < 0.0001; ns, not significant.

### IFNγ production by CD4 T cells is largely redundant at the FRT mucosa

The complete lack of IFNγ production in IFNγ^−/−^ mice only allows the evaluation of the global effect of this cytokine that derives from numerous cellular sources. To specifically define the contribution of CD4 T cell-dependent IFNγ production to host resistance to *Chlamydia*, we conducted T cell adoptive transfer experiments in which we transferred either IFNγ-sufficient (WT) or IFNγ-deficient (IFNγ^−/−^) CD4 T cells into the innate immunity intact TCRβ^−/−^ mice, and challenged the recipients intravaginally with *C. muridarum* (Fig. 4A). Compared to WT CD4 T cell transfer group, TCRβ^−/−^ mice receiving IFNγ-deficient CD4 T cells exhibited similar rates of bacterial containment for the first 21 days (fold change in log_10_: 3.1±1.8 in WT CD4 transfer vs 2.7±1.2 in IFNγ^−/−^ CD4 transfer, p=0.21; Fig. 4B). *Chlamydia* burden continued to decrease for another 10-fold in the IFNγ^−/−^ CD4 T cell transfer group before these mice entered the chronic, low-grade shedding phase around day 35 (Fig. 4B). The accumulative ~5,000-fold decrease in bacterial burden demonstrated that IFNγ^−/−^ produced by CD4 T cells is dispensable for eliminating vast majority of the pathogen from the FRT. Lastly, as a result of the functional innate immunity in TCRβ^−/−^ recipient mice, the lethal dissemination phenotype was completely rescued in this adoptive transfer model compared to IFNγ^−/−^ mice, reinforcing the idea that innate IFNγ is sufficient for systemic *Chlamydia* containment (Fig. 4C).

**Fig. 4.**
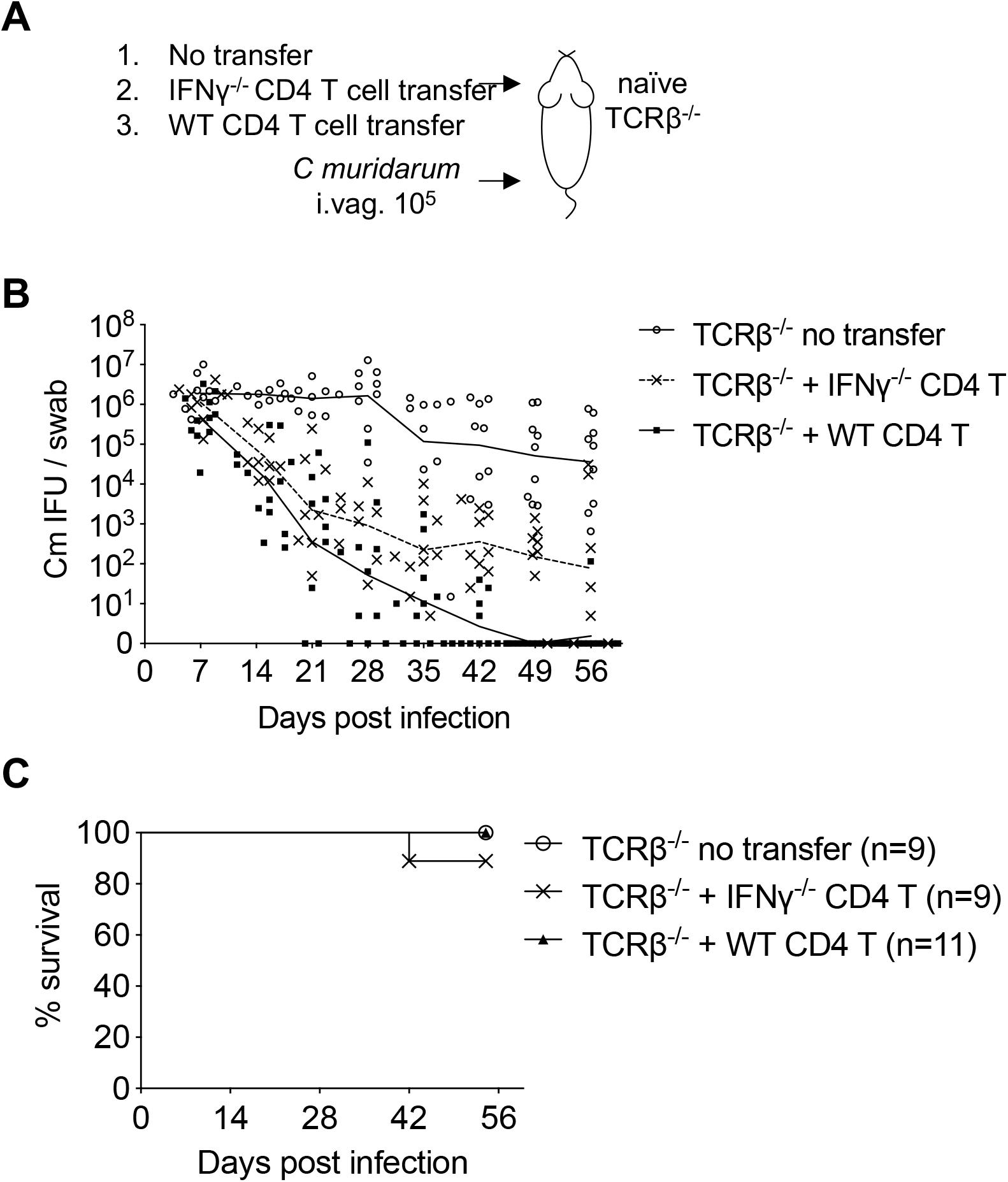
CD4 T cell-dependent IFNγ production is largely redundant at the FRT mucosa. CD4 T cells isolated from WT B6 or IFNγ^−/−^ mice were adoptively transferred to TCRβ^−/−^ mice. Recipient TCRβ^−/−^ mice were infected intravaginally with 1×10^5^ *C. muridarum*. (A) Schematic depicting the TCRβ^−/−^ adoptive transfer experimental setup. (B) Bacterial shedding from the FRT and (C) survival of TCRβ^−/−^ recipient mice after receiving naïve CD4 T cells from either WT or IFNγ^−/−^ donors and intravaginal infection. Data shown are combined results of three independent experiments of 6-11 mice per group. Each data point in (B) and (C) represents an individual mouse. Lines represent mean values.

## Discussion

The ability of T cells to produce IFNγ in response to infection is an important readout for their antigen specificity and effector function. Although Th1 cells are essential for combating intracellular parasites, the reliance on IFNγ to confer protection appears to be pathogen-specific. Mice lacking IFNγ receptor or Th1 transcription factor T-bet are highly susceptible to *Salmonella* infection (26, 27). In contrast, effective control of *Mycobacteria tuberculosis* infection depends on Th1 cells but has a minimal requirement of IFNγ in the lung (28–30). The importance of CD4 T cells in host resistance to *Chlamydia* is highlighted by the absolute requirement of MHCII and αβTCR for bacterial control in the FRT mucosa, while the most prominent phenotype of IFNγ^−/−^ mice is lethal bacterial dissemination (14, 15, 24). By conducting side-by-side comparison of TCRβ^−/−^ and IFNγ^−/−^ mice in this current study, we confirmed previous findings and argued unambiguously that hosts had distinct requirements for T cells and IFNγ for *Chlamydia* resistance.

TCRβ^−/−^ mice lack αβT cells but retain relatively intact innate immunity and several adaptive immune components such as γδT cells and T-independent antibody, all of which may contribute to the long-term survival of the hosts. As part of the innate immune system, ILCs are known for their functions that parallel with their T cell counterparts, but differ from T cells by their embryonic origin, tissue-residency, lack of rearranged receptor for Ag recognition and unique contribution to tissue integrity (31). Immune responses of the ILCs are critical for early control of pathogen replication before the host launches an effective adaptive immune response. This defense mechanism could be particularly important for containing mucosal pathogens like *Chlamydia*, since the target FRT tissue lacks defined lymphoid structure and consequently effective adaptive immune responses are significantly delayed (32, 33). Using the *Rag1*^−/−^ and *Rag2*^−/−^*γc*^−/−^ models, we show in this current study that removing ILCs from the innate immune system had a detrimental effect on host resistance to *Chlamydia*, as *Rag2*^−/−^*γc*^−/−^ mice quickly succumb to disseminated infection with high-grade bacteremia. By contrast, ILC-sufficient *Rag1*^−/−^ mice exhibited significantly longer survival and stable body weights. These observations made it evident that ILCs are essential for preventing early systemic *Chlamydia* dissemination in the absence of adaptive immunity. Given the early responses of ILCs, it is unlikely that ILCs are completely redundant to adaptive responses to prevent early bacterial dissemination in immunocompetent hosts (34). This assumption needs to be confirmed experimentally by future studies.

Our IFNγ and NK1.1 depletion experiment addressed directly that both innate IFNγ and NK1.1^+^ group 1 ILCs contribute to host resistance to lethal *Chlamydia* dissemination in *Rag1*^−/−^ mice. Group I ILCs, including cytotoxic NK cells and non-cytotoxic ILC1s, are the major source of IFNγ during innate immune responses along with neutrophils, macrophages, DCs and a subset of ILC3s (35–37). In immunocompetent hosts, IFNγ production by NK cells can be detected as early as 4 hours after intravaginal *C. muridarum* infection (38). The early response of NK cells dictates CD4 T cell differentiation and memory responses during *C. muridarum* lung infections (39, 40). Recently, Poston et al. have shown that T cell-independent IFNγ cooperates with B cells to prevent lethal dissemination caused by a highly virulent *C. muridarum* strain CM001 (17). Our data add to previous findings by showing ex vivo IFNγ secretion on the surface of CD11b^+^ NK1.1^+^ cells, indicating that group 1 ILCs elicit protective function, as least in part, by their production of IFNγ in the absence of T cells. It should be noted that both IFNγ-depleted and NK1.1-depleted *Rag1*^−/−^ mice exhibited slightly longer survival times than *Rag2*^−/−^*γc*^−/−^ mice, indicating that additional cellular sources, such as ILC3s, may also produce IFNγ and confer protection against *Chlamydia* as shown in the mouse endometrium tissue in a recent study (41). Additionally, other protective mechanisms independent of IFNγ may be involved in ILC-mediated protection. It is intriguing to observe that serum IFNγ was not significantly affected by anti-NK1.1 treatment in *Rag1*^−/−^ mice and was minimally detected in WT animals. These results suggest two non-mutually exclusive probabilities: first, circulating IFNγ is neither necessary nor sufficient for protection. Instead, close proximity between IFNγ secreting cells and responders are essential for bacterial control (42). Second, circulating IFNγ is more efficiently removed by IFNγ receptor-expressing cells in immune-sufficient mice (43). Future experiments are required to address these issues directly.

The sustained *Chlamydia* replication in the FRT of TCRβ^−/−^ mice suggested that innate immune cells, γδT cells, and T-independent Abs are all incapable of restraining *Chlamydia* growth in the FRT. In contrast, adoptive transfer of CD4 T cells, regardless of their ability of producing IFNγ, reduced bacterial burden to less than 0.1% of the peak burden within 3 weeks. Our results are in parallel with previous reports of IFNγ^−/−^ mice but differ from those by retaining intact innate IFNγ in our system thereby lethal disseminated infection was prevented (14, 24). By only manipulating the CD4 T cell compartment, we confirmed that CD4 T cells are both necessary and sufficient for *C. muridarum* control in the FRT. More importantly, we shown unequivocally that CD4 T cell-derived IFNγ is largely redundant for protective immunity in the FRT, at least during the early phase of bacterial replication.

Numerous studies in the field have firmly established the importance for the pleiotropic cytokine IFNγ in *C. muridarum* and *C. trachomatis* infection, in vitro and in vivo (44–55). Indeed, our data along with many others have demonstrated that complete eradiation of *C. muridarum* from the FRT relies on CD4-derived IFNγ (14, 24). Nevertheless, our study emphasized that a highly effective mechanism of T helper cell-mediated protection against mucosal *Chlamydia* infection is yet to be discovered. Efforts in searching for such protective mechanism is urgently needed as precise understanding of protective immunity is fundamental to the rational design of a much-needed Chlamydia vaccine. Fortunately, recent studies started to shed light on several important aspects of CD4 T cell biology related for host protective responses beyond IFNγ production. Yu et al. showed that vaccination using live or dead *C. muridarum* EB elicits different degrees of protection that correlates with the frequency of multifunction Th1 cells (20). Likewise, a recently developed *C. muridarum* TCR-transgenic model has revealed that the ability of monoclonal Ag-specific CD4 T cells to co-produce IFNγ, TNFα and IL-2 is essential for their protective efficacy (56). Using MHC class II tetramers, we demonstrated that *C. muridarum* infection induces a highly heterogenous T helper response dominated by Th1 while accompanied by fractions of *Chlamydia*-specific Treg and Th17 cells in lymphoid and mucosa tissues (33). Other Th lineages, such as an IFNγ IL-13 double producing CD4 T cell clone, and a Th2-dominant response in human *C. trachomatis* infection have also been documented (57, 58). Finally, an elegant *C. trachomatis* vaccine study conducted by Stary et al. has revealed that T cell activation, effector functions and formation of tissue-resident memory should all be taking into consideration when evaluating protective efficacy afforded by a vaccine (59). Our knowledge of CD4 T cell differentiation is evolving quickly as a result of groundbreaking technologies. It is proposed that a continuum of CD4 T cell states will likely replace our traditional understanding of defined CD4 T cell lineages (60). With the increased knowledge and higher resolution tool to understand CD4 T cell biology, a refreshed notion of protective immunity against *Chlamydia* will likely emerge.

## Acknowledgements

We thank Dr. Richard Morrison and Sandra Morrison for providing us the PK136 monoclonal antibody. We thank Andrea Harris at the UAMS Flow Cytometry Core for technical assistance. This study was supported by grants from the National Institutes of Health to LXL (AI139124 and GM103625).

